# Overt social interaction and resting state in young adult males with autism: core and contextual neural features

**DOI:** 10.1101/332213

**Authors:** Kyle Jasmin, Stephen J. Gotts, Y. Xu, S. Liu, Cameron Riddell, John Ingeholm, Lauren Kenworthy, Gregory L. Wallace, Allen R. Braun, Alex Martin

## Abstract

Conversation is an important and ubiquitous social behavior. Individuals with Autism Spectrum Disorder (autism) without intellectual disability often have normal structural language abilities but deficits in social aspects of communication like pragmatics, prosody, and eye contact. Previous studies of resting state activity suggest that intrinsic connections among neural circuits involved with social processing are disrupted in autism, but to date no neuroimaging study has examined neural activity during the most commonplace yet challenging social task: spontaneous conversation. Here we used functional MRI to scan autistic males (N=19) without intellectual disability and age- and IQ-matched typically developing controls (N=20) while they engaged in a total of 193 face-to-face interactions. Participants completed two kinds of tasks: Conversation, which had high social demand, and Repetition, which had low social demand. Autistic individuals showed abnormally increased task-driven inter-regional temporal correlation relative to controls, especially among social processing regions and during high social demand. Furthermore, these increased correlations were associated with parent ratings of participants’ social impairments. These results were then compared with previously-acquired resting-state data (56 Autism, 62 Control participants). While some inter-regional correlation levels varied by task or rest context, others were strikingly similar across both task and rest, namely increased correlation among the thalamus, dorsal and ventral striatum, somatomotor, temporal and prefrontal cortex in the autistic individuals, relative to the control groups. These results suggest a basic distinction. Autistic cortico-cortical interactions vary by context, tending to increase relative to controls during Task and decrease during Rest. In contrast, striato- and thalamocortical relationships with socially engaged brain regions are increased in both Task and Rest, and may be core to the condition of autism.

## Introduction

Conversation is an important part of everyday life that provides a means to share information and express affiliation. Abnormal brain development during childhood is associated with conditions in which people struggle with conversation skills, as in the case of Autism Spectrum Disorder (referred to hereafter as “autism”). This class of neurodevelopmental conditions is characterized by deficits in social functioning, communication, repetitive behaviors, and stereotyped interests (Lord *et al.*, 2000). Structural language *per se* may be unimpaired except where it intersects with social processing (Mundy *et al.*, 1990) such as in pragmatics (Baron-Cohen, 1988; Loukusa and Moilanen, 2009), and prosody (Paul *et al.*, 2005; Shriberg *et al.*, 2001). Non-verbal aspects of communication such as eye gaze (C. D. Frith and U. Frith, 1999; Loveland and Landry, 1986; Pelphrey *et al.*, 2005) and hand gestures (Attwood *et al.*, 1988; de Marchena and Eigsti, 2010) are also often impaired, and elicitation of deficits in these behaviors through tools like the ADOS (Autism Diagnostic Observation Schedule, (Lord *et al.*, 2000) is standard diagnostic procedure. Indeed, one of the DSM-5 (Diagnostic and Statistical Manual of Mental Disorders) Social-Communication symptoms of autism is problems with social-emotional reciprocity as seen in normal conversational turn-taking. Severe impairments in social interaction have been linked with difficulty establishing and maintaining social relationships (Orsmond *et al.*, 2004), thus abnormal conversation behavior can have serious consequences for people with autism.

Despite conversation’s ubiquity and social importance, no published neuroimaging studies have measured neural activity in autistic individuals while they are engaged in face-to-face conversation with naturalistic sensory, cognitive, communicative and social demands. Instead, studies have focused on the components of conversation with which people with autism struggle. Investigations of vocal prosody have found abnormally increased brain activity both within the same network of regions active in typically developing control participants (Wang *et al.*, 2006) and across diffuse regions outside of typical networks (Colich *et al.*, 2012; Eigsti *et al.*, 2012). Studies of pragmatic processing have found that integrating linguistic and real-world knowledge is associated with abnormally low levels of activity in left IFG in autistic children (Groen *et al.*, 2010) but abnormally high levels in homologous right IFG in autistic adults (Tesink *et al.*, 2009).

Meanwhile, resting-state studies evaluating intrinsic brain organization have indicated that disordered connections between cortical regions may play a role. The bulk of the resting-state literature on autism has focused on high-functioning adolescent and adult males using functional MRI (see Hull *et al.*, 2017; Picci *et al.*, 2016 for reviews). In seed-based connectivity studies using a small number of seeds, researchers have reported a variety of differing patterns of functional connectivity, some that show decreases in autism among cortical regions (e.g. Abrams *et al.*, 2013; Alaerts *et al.*, 2014; Ebisch *et al.*, 2011; Fishman *et al.*, 2015; Jung *et al.*, 2014; Kennedy and Courchesne, 2008; Linke *et al.*, 2018; Verly *et al.*, 2014; Weng *et al.*, 2010) and some that show increases (e.g. Alaerts *et al.*, 2014; Chien *et al.*, 2015; Fishman *et al.*, 2014; 2015; Nebel *et al.*, 2014a, 2014b; Redcay *et al.*, 2013). Among these, those investigating seeds used to define the so-called "default mode" network (e.g. Raichle, 2015), sharing some regions with the language system (e.g. Turken and Dronkers, 2011; Jasmin *et al.*, 2016; Scott *et al.*, 2000), have often found decreased functional connectivity among related regions, such as the posterior cingulate, medial prefrontal cortex, and the parahippocampal gyrus (e.g. Assaf *et al.*, 2010; Kennedy and Courchesne, 2008; see also Starck *et al.*, 2013; Weng *et al.*, 2010; Ypma *et al.*, 2016). Other studies have attempted to simultaneously examine all possible combinations of regions (so-called "data-driven" studies), with several of these showing decreased functional connectivity amongst cortical regions associated with social functions, such as the STS, medial prefrontal, temporoparietal junction, left IFG, as well as somatosensory cortex (e.g. Anderson *et al.*, 2011; Cerliani *et al.*, 2015; Cheng *et al.*, 2015; Di Martino *et al.*, 2014; Gotts *et al.*, 2012; von dem Hagen *et al.*, 2013), with some showing simultaneously increased functional connectivity between thalamus, striatum and some of the same cortical regions (e.g. Cheng *et al.*, 2015; Cerliani *et al.*, 2015; Di Martino *et al.*, 2014, Linke *et al.*, 2018; see also Delmonte *et al.*, 2013; Di Martino, Ross, *et al.*, 2009; Nair *et al.*, 2015; Padmanabhan *et al.*, 2013; although see Nair *et al.,* 2013). For cohorts that also include even younger participants, increased cortico-cortical functional connectivity has been observed (e.g. Abbott *et al.*, 2016; Supekar *et al.*, 2013; Uddin *et al.,* 2013; although see also Dinstein *et al.*, 2011). There is also some question as to whether females with autism exhibit the same patterns of functional connectivity alterations as males (e.g. Alaerts *et al.*, 2016; Floris and Howells, 2018; Lai *et al.*, 2017; Ypma *et al.*, 2016). Other studies have highlighted additional and complex aspects of the resting-state functional connectivity changes (Hahamy *et al.*, 2015).

Here we investigate the neural basis of face-to-face verbal interactions in matched samples of adolescent and adult males with and without an autism diagnosis. Participants were scanned with functional MRI while they spoke via cameras and microphones with a real interaction partner. The interactions took place under two conditions: *Conversation* – wherein the participant spoke spontaneously and reciprocally with their partner, and *Repetition* – in which the subject merely repeated their partner’s speech. These two tasks had similar sensory and motor components (both involving turn-taking), but differed crucially in their level of social demand – Conversations (but not Repetitions) involved novel, spontaneous and reciprocal speech that relied on social knowledge.

We had two main objectives. First, we used a whole-brain approach to evaluate changes in between-region correlations in our two groups (Autism vs Control) and tasks (Conversation vs Repetition). Then we compared group differences in inter-region correlations at task with a larger sample of resting state data, to identify commonalities and differences between the two states.

## Materials and Methods

### Participants

Nineteen (19) males (aged 14.7 - 28.2 years) with autism and 20 male control participants (aged 15.1 to 32.0 years) took part in the tasks. Participants with autism were recruited from the Washington, DC, metropolitan area and met DSM-5 criteria for ASD (APA, 2013) as assessed by an experienced clinician. All participants with autism received the ADOS module 4 (Lord *et al.*, 2000). The scores from participants with autism met cut-off for the ‘broad autism spectrum disorders’ category according to criteria established by the National Institute of Child Health and Human Development/National Institute on Deafness and Other Communication Disorders Collaborative Programs for Excellence in Autism (Lainhart *et al.*, 2006). Scores on the Social Responsiveness Scale (Constantino *et al.*, 2003), a measure of social impairment, were obtained by parent-report. The distributions for full-scale IQ, verbal IQ, and age did not differ significantly between the autism and control groups (Supplementary Table 1).

### Procedure

Each session consisted of five functional MRI runs. Runs 1, 3 and 5 were Conversations and runs 2 and 4 were Repetition. Prior to scanning, participants were told that they would engage in unstructured and informal conversations with the experimenter. Using a modified version of the Interest Scale questionnaire (Anthony *et al.*, 2013; Bodfish, 2003), participants rated their level of interest in various topics such as "music", "games", and "transportation vehicles", and indicated their top three interests, from which the experimenter selected two. The topic of the final conversation was always work or school life, depending on participant age. The topics of conversations were coded to match the categories on the Interests Scale and are listed in Supplementary Table 2. Participants chose the two nursery rhymes they would recite from a list they received before the session.

Before each run, the experimenter sat in front of a blue screen facing a camera. The run began with 16 seconds of rest (8 TRs), during which the participant saw the word "REST". Then, live video and audio from the experimenter were presented to the subject and the verbal interaction began (see Supplemental Methods). The experimenter always initiated the interaction. Conversations proceeded for 6 minutes, and Repetitions for 3 minutes. After each interaction, the video faded to black and a "STOP" slide was displayed to the participant, followed by 30 additional seconds (15 TRs) of rest to allow for delayed hemodynamic effects. The study was designed such that participants spent more time engaging in Conversation than Repetition in order to limit the duration of the entire session as a whole, and to ensure participants did not become fatigued during Repetition runs.

### MRI data acquisition

T2*-weighted BOLD images were acquired on a General Electric Signa HDxt 3.0 Tesla scanner (GE Healthcare, Waukesha, WI, USA) with an 8-channel head coil. A single-shot gradient-echo echo planar imaging sequence was used: the acceleration factor of ASSET (Array Spatial Sensitivity Encoding Technique) = 2, TR = 2000 ms, TE (echo time) = 30 ms, flip-angle = 90°, 64×64 matrix, field of view = 227 mm. 40 interleaved sagittal slices with a thickness of 4 mm were used to cover whole brain (voxel size = 3.55 x 3.55 x 4 mm^3^). Sagittal acquisition was used to help minimize slicing artifact that can occur when using more typical axial acquisitions during overt speech (e.g. Birn *et al.*, 2010). The Conversation runs consisted of 203 TRs and the Repetition runs consisted of 113 TRs. A high-resolution T1-weighted anatomical image (MPRAGE, Magnetization-Prepared Rapid Gradient-Echo) was also obtained (124 axial slices, 1.2 mm slice thickness, field of view = 24 cm, 224 x 224 acquisition matrix).

### MRI data pre-processing

Echo-planar image preprocessing was performed with AFNI (Cox, 1996). The first four TRs of each run were removed to allow for T1 stabilization, and outlying time points in each voxel, corresponding to motion and other artifacts, were attenuated with AFNI’s 3dDespike. Volumes were slice-time corrected and co-registered to the anatomical image. Sources of motion-related and physiological noise were removed with ANATICOR (Jo *et al.*, 2010; 2013) and aCompCorr (Behzadi *et al.*, 2007). The regressors for these methods were created by first segmenting the anatomical scan into tissue types with Freesurfer (Fischl *et al.*, 2002). An eroded ventricle mask was applied to the volume-registered echo-planar image data to create a nuisance time series for the ventricles. An eroded white matter mask was used to create a "localized" estimate of the BOLD signal in white matter, which was averaged within a 20-mm radius sphere centered on each voxel (see Jo *et al.*, 2010; 2013 for further discussion). For runs with physiological data, 8 Retroicor (Glover *et al.*, 2000) and 5 respiration volume per time (Birn *et al.*, 2008) regressors were created from the cardiac and respiration measures, estimated at slice time 0. The multiple physiological regressors of each type represent several interpolated time points within each TR. Additional speech-related motion and respiration artifacts were modeled using a PCA decomposition of BOLD fluctuations in non-gray-matter locations (Behzadi *et al.*, 2007). Eroded masks of white matter and ventricles were joined into a single nuisance tissue mask and applied to each re-aligned and co-registered functional time series, which were detrended by 4th order polynomials prior to PCA in order to remove scanner drift and other very low frequency signals. The time series were then decomposed with PCA, retaining the first 3 principal components, as pilot analyses indicated these were the components that contained the most variance related to model-based respiration regressors (Retroicor and RVT regressors; see also Stoddard *et al.*, 2016). The full nuisance regression model for each voxel therefore included regressors for the 4th order polynomial baseline model, 1 ventricle time series, 1 localized white-matter time series, 6 motion parameters, 5 respiration volume per time regressors, 8 Retroicor time series (4 cardiac and 4 respiration regressors) and 3 aCompCor principal component regressors. The “clean” residual time series of this model was converted into Talairach space, resampled to 3mm^3^ isotropic voxels, and used in all subsequent analyses.

The magnitude of transient head motion was calculated from the 6 motion parameters and aggregated as a single variable using AFNI’s @1dDiffMag to calculate a Motion Index (Berman *et al.*, 2016; Gotts *et al.*, 2012). This measure is comparable to average Framewise Displacement over a scan (Power *et al.*, 2012) and is in units of mm/TR. The grand average correlation of all voxel time series with each other, or GCOR (for global correlation level), was also calculated for each experimental run to serve as a more omnibus measure of residual global artifacts and was used as a nuisance covariate in ROI-based, group-level analyses (Gotts *et al.*, 2017; Gotts, Jo, *et al.*, 2013; Saad *et al.*, 2013; Zachariou *et al.*, 2017).

### Task-related functional connectivity analyses

Our main analyses utilized a functional connectivity approach in which correlations among voxel- and/or ROI-based time series were compared across conditions. In part, this choice was mandated by the lack of appropriately spaced baseline periods during naturalistic conversation, preventing the use of a more typical general linear model approach. This form of analysis was also directly comparable to that applied previously to resting-state data. Our experiment used a 2×2 mixed factorial design with one within-subjects variable (Task) and one between-subjects variable (Group). Two types of relationships were identified: 1) inter-region correlations that differed between the autism and control groups regardless of task (i.e. main effect of Group); and 2) inter-region correlations that differed by Group and Task (their interaction). Our functional connectivity analyses employed the 3 steps used by Gotts and colleagues (see also Berman *et al.*, 2016; Gotts *et al.*, 2012): seed definition, target ROI selection, and region-to-region correlation analysis. We undertook all 3 steps for both the Group effect and the Group by Task interaction effects. For completeness, we also report the main effect of Task (Conversation>Repetition).

Seeds were identified using whole-brain "connectedness" (e.g. Cole *et al.*, 2010; Gotts *et al.*, 2012). The average Pearson correlation of each voxel’s time series with every other voxel’s time series was calculated to create a 3D reduction of the 4D (3D + Time) dataset for each functional run (available as the AFNI function 3dTcorrMap). Since connectedness reflects the average level of correlation with the rest of the brain, it gives an indication of how involved a given brain area was with the task during the scan. This approach, akin to *centrality* in graph theory, has been used previously in studies of resting-state (Berman *et al.*, 2016; Cole *et al.*, 2010; e.g. Gotts *et al.*, 2012; Meoded *et al.*, 2015; Stoddard *et al.*, 2016; Watsky *et al.*, 2018) and task-based functional connectivity (Song *et al.*, 2015; Steel *et al.*, 2016). Linear mixed effects models (Chen *et al.*, 2013) were constructed whose dependent variables were the voxel-wise connectedness maps from each functional run. Group and Task and their interaction were included as a fixed effects. Participant Age and the Motion Index (computed separately for each run) were included as nuisance covariates. Participant was treated as a random intercept. Cluster correction was used to control the Type I error rate. The average smoothness of the cleaned functional time series was estimated with AFNI’s 3dFWHMx, using the empirical, spatial autocorrelation function (June 2016). 3dClustSim (June 2016) was then used to run a Monte Carlo simulation with 5,000 iterations within the gray matter mask in Talairach space that the analyses were performed within. Importantly, the smoothness estimates and noise simulations did not assume Gaussian distributions of activity, which has been shown to inflate the false positive rate in studies using more traditional cluster size correction (Cox *et al.*, 2017; e.g. Eklund *et al.*, 2016). Clusters were selected at a cluster defining threshold of p<.001, minimum cluster size k=22.

The seed definition step was then followed with more typical seed-based correlation analyses. Signal within each seed region was averaged across voxels to form ROI-averaged time series, which were correlated with the time series for every voxel in the brain, separately for each run. These correlations were Fisher z-transformed and used as dependent variables in linear mixed effects models with the same fixed and random effects as the previous step. We tested for the main effect of Group at a voxel threshold of *p*<.001, with correction by cluster size for whole-brain comparisons as well as the number of seeds tested (i.e. FWE correction to *p*<[.05 / (number of seeds)]. Results of the separate seed-based tests were combined to form one composite map, first by binarizing each seed-to-whole-brain test into zeros and ones and then summing across them. This map was then thresholded at 80% of the maximum possible sum (i.e. if 6 seeds were tested, included voxels would need to have arisen in 5/6 tests). Clusters smaller than 20 voxels were then excluded in order to eliminate small singleton clusters with higher levels of voxel noise. Secondary target regions were then combined together with seed regions to arrive at a full set of ROIs. Region-by-region matrix analyses were then conducted using the same contrasts applied to connectedness and the seed-based tests, allowing the examination of all inter-regional relationships. For these all-to-all matrix tests, as well as correlation with social impairment and comparison with resting state data, GCOR was included as an additional nuisance covariate (to Age and Motion Index) in order to insure that any group effects were not due to due to residual global or speech-related artifacts (Gotts *et al.*, 2017; Gotts, Jo, *et al.*, 2013; Saad *et al.*, 2013).

Correlations with social impairment as measured with the Social Responsiveness Scale (SRS-1 Total Raw Score, Constantino *et al.*, 2003; Gotts *et al.*, 2012; Ramot *et al.*, 2017) were tested with additional models in the autism subjects, separately for the Group Main Effect ROIs and the Group X Task Interaction ROIs. The analyses of Group Main Effect ROIs included data from both tasks, Conversation and Repetition, whereas the analysis of Group X Task Interaction ROIs focused on the Conversation runs, which were of primary interest.

Multiple comparisons in the region-by-region correlation tests were controlled with False Discovery Rate (e.g. Genovese *et al.*, 2002), calculated by pooling over *p*-values resulting from task matrix tests together (autism vs. control for the Group ROIs; Interaction test in Group X Condition ROIs; effects of SRS in both of these matrices). We report all results un-thresholded in the lower triangles of the matrices, with corrected results (*q*<.05) in the upper triangles.

### Comparison of task and resting state data

Several previous studies of functional connectivity at rest in male adolescents and adults with autism have observed a pattern of cortico-cortical decreases among several brain regions engaged by social tasks, such as the STS, medial prefrontal cortex, the temporoparietal junction, left IFG, and somatosensory cortex (e.g. Anderson *et al.*, 2011; Cerliani et al., 2015; Cheng *et al.*, 2015; Di Martino *et al.*, 2014; Gotts *et al.*, 2012; see Picci *et al.*, 2016, Hull *et al.* 2017 for reviews). We included an explicit comparison of results in task and rest in the current study to determine which patterns of functional connectivity differences are robust across cognitive states with and without overt motor behaviors versus those that are context-specific to state. The resting state data we included have been used in previous publications (Gotts *et al.*, 2012; 2017; Gotts, Jo, *et al.*, 2013; Gotts, Saad, *et al.*, 2013; Plitt *et al.*, 2015; Power *et al.*, 2017; Ramot *et al.*, 2017) and were collected on the same 3 Tesla MRI scanner as the task data. The preprocessing for these data also used the same ANATICOR procedure as for the task data with the only difference being the lack of the aCompCor regressors (Behzadi *et al.*, 2007). For rest scans, participants were instructed to lie still and relax, maintaining fixation on a central cross on the viewing screen. Participants included 56 autistic males (mean age = 19.1 years, SD = 3.8 years) and 62 control males (mean age = 21.2 years, SD = 5.1 years), with the same selection/inclusion criteria as discussed above for the task data. Participant groups were matched on age, IQ, head motion, and overall measures of temporal signal to noise ratio. Informed assent and consent were obtained from all participants and/or their parent/guardian (participants younger than 18), and the experiment was approved by the NIMH Institutional Review Board (protocol 10-M-0027, clinical trials number NCT01031407).

Two sets of analyses were conducted with resting state data. In the first, the regions identified as showing a main effect of group during task were applied directly to the resting-state data for quantitative comparison with task using linear mixed effect models (AFNI’s 3dLME), with Conversation and Repetition data pooled to form the condition ‘Task’. Each region-by-region combination of functional connectivity served as dependent variables, with Group and State (Task, Rest) included as fixed effects. Main effects of Group were evaluated within Task and Rest separately, along with a Group X State interaction (correction for multiple comparisons by False Discovery Rate to *q*<.05). The second analysis compared whole-brain connectedness between autism and control participants in Rest, with Age and Motion Index as nuisance covariates for comparison with the main effect of Group during Task in terms of spatial overlap (voxelwise threshold of p<.001, FWE corrected using cluster size to *p*<.05; at this voxelwise threshold, results were also corrected by False Discovery Rate to *q*<.05).

## Results

### Verbal output

First we examined whether participants in the autism and control groups performed the task similarly, to ensure any group differences in the neuroimaging analysis could not be explained by gross differences in task-relevant behavior. Audio recordings were analyzed with MATLAB and raw counts of words and sentences were obtained following transcription by a professional transcription service (see Supplement). We examined SpeakingTime (defined below), total number of words uttered, number of speaking turns taken, and number of words per sentence.

For each run, SpeakingTime was calculated as the ratio of time a participant spoke during a run divided by the total time spent speaking by both the participant and experimenter [PPTspeech / (PPTspeech + EXPspeech)]. Ratios greater than 0.5 indicated that the participant talked more than the experimenter. Overall there were no differences between groups or conditions for the Conversation (t(37)=0.31, p=.76) or Repetition runs (t(37)=0.66, p=0.51. Supplementary Fig. 1). The total number of words produced also did not differ between the autism and control groups (t(37) = −1.4, p=0.17), and neither did the number of speaking turns t(37) = 0.46, p= 0.65), nor the number of words per sentence t(37) = 0.01, p = 0.99). For the repetition runs, the participant repeated exactly what their partner spoke, resulting in exactly matched verbal output.

### Motion

Motion Index values for all runs individually (Supplementary Fig. 2) were subjected to a 2×2 repeated measures ANOVA with Group, Task, and Group-by-Task as terms. Motion did not differ by Task (F(1,74) = 0.92, p=.34), and Task and Group did not interact to predict motion (F(1,74) = 0.47, p=0.50). However, there was a statistically significant main effect of Group (F(1,74) =9.1, p=.003). To assess the overall magnitude of this group difference, we calculated the average Motion Index by subject and by group, and compared them. The ASD group moved 57 microns per TR (about 1/17^th^ of a millimeter) more than the TD controls (Mean ASD Motion = 0.216±.09mm/TR. Mean TD motion = 0.159±.07mm/TR). To mitigate any remaining motion-related artifacts that were not already removed by our cleaning procedure, the Motion Index for each run was included as a covariate in all subsequent correlation analyses. A control analysis excluding high motion runs in the Autism group and low motion runs in the Control group in order to match motion levels by group also did not lead to large changes in the results reported below.

### Comparison of Conversation vs. Repetition in whole-brain connectedness

Due to the limitations of a general linear model approach during naturalistic conversation without appropriately spaced baseline periods, we instead took a basic correlational (task-driven functional connectivity) approach to the main analyses of interest (although see Supplementary Fig. 3 for a restricted general linear model analysis of local activity differences in Speaking versus Listening periods). Using this approach, we first examined the Main Effect of Task on whole-brain “connectedness” (the correlation of each voxel with the rest of the brain; Cole *et al.*, 2010; Gotts *et al.*, 2012; Salomon *et al.*, 2011). The main effect of Task yielded highly significant results with multiple large clusters (voxelwise threshold of *p*<.001). The strongest differences in connectedness (Conversation > Repetition) were observed in in dorsomedial prefrontal cortex, left fronto-temporal cortex with a peak in left anterior temporal lobe (prominently including the pole and STG), and left angular gyrus (Supplementary Table 3; Fig. 1).

**Figure 1:**
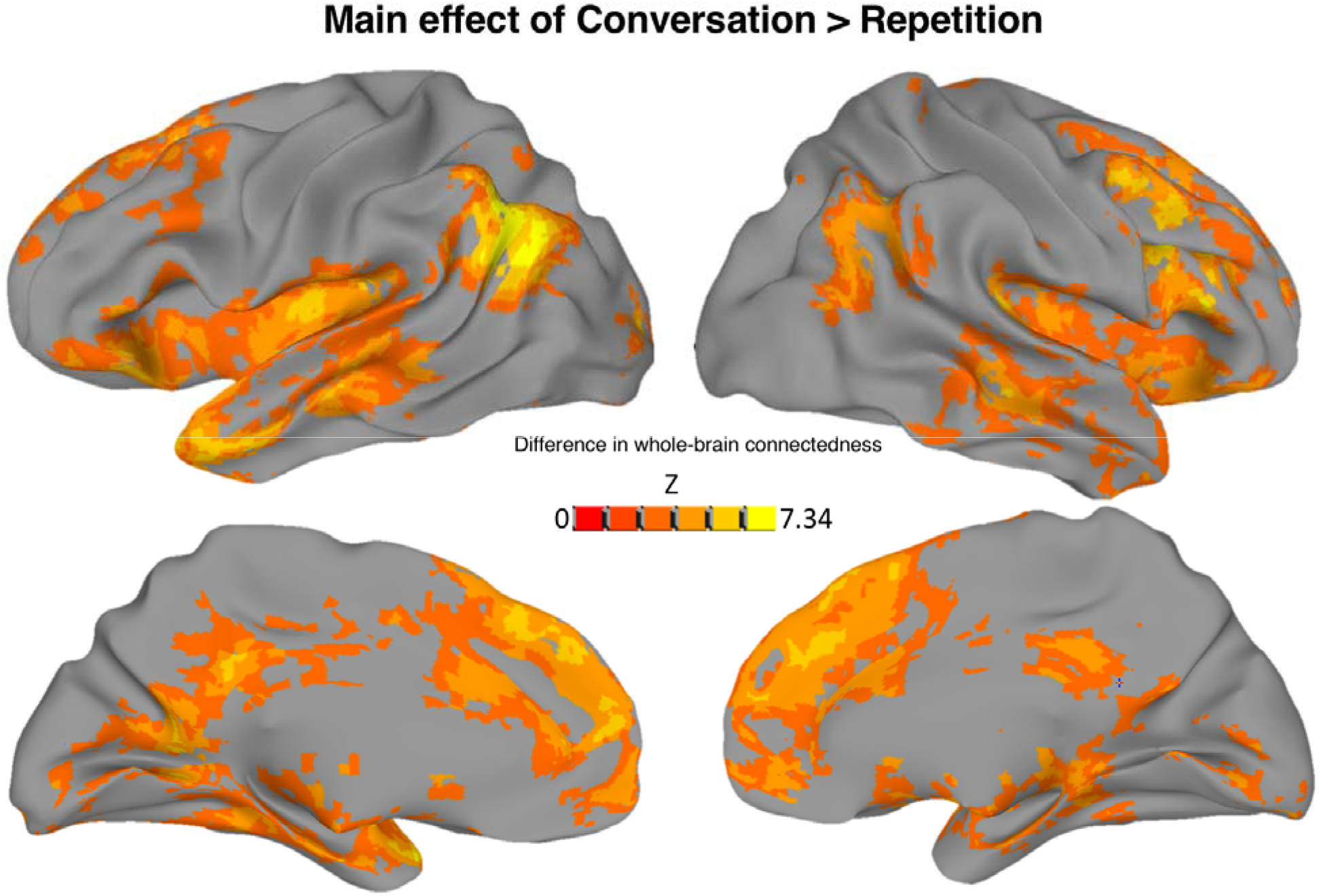
Conversation vs Repetition. A test for the Main effect of Task, across all whole-brain connectedness values from both groups, revealed a number of areas that showed greater whole-brain involvement during Conversation compared with the Repetition condition. The strongest of these results were frontal and temporal areas (especially left temporal pole), the left temporoparietal junction, and bilateral dorsomedial prefrontal cortex. Voxels significant at p<.001 are plotted on an inflated Freesurfer standard surface (Fischl *et al.*, 2002).

### Main effect of Group

The main effect of Group on connectedness values identified 6 regions whose whole-brain connectedness was greater in the autism than the control group. There were no regions showing greater correlation for the control than the autism group. Increased correlation for the autism participants was found for two clusters in right anterior temporal lobe, a cluster at the left temporal pole, the right inferior frontal gyrus and anterior insula, right ventral striatum, and somatomotor cortex (Supplementary Table 4). Using these clusters as seeds in a seed-to-whole-brain analyses (see Methods), we identified 10 additional regions that showed greater correlation in autism than control participants (Fig. 2; Supplementary Table 5). An analysis of the correlation values within the region-by-region matrix revealed a number of regional pairs whose correlations differed between the autism and control groups. This was most pronounced among right hemisphere regions such as the precuneus, anterior superior temporal gyrus and sulcus, and IFG. The most implicated region was a cluster in the right ventral striatum, which showed a significant hyper-correlation with each of the other regions, even after multiple comparisons correction (Fig. 2).

**Figure 2:**
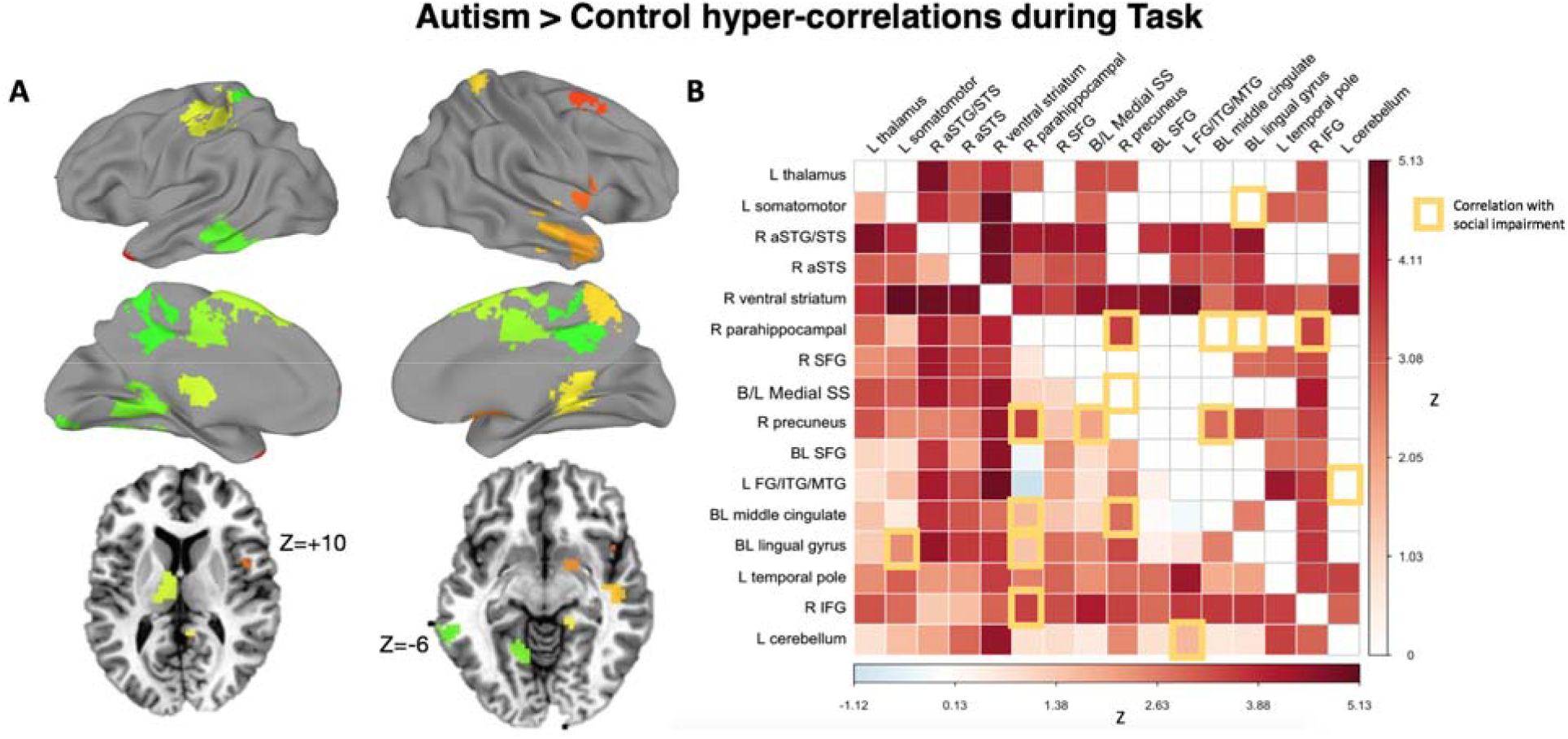
Group (Autism > Control) comparison of inter-region correlations across tasks. **A)** Regions of interest defined by the Main Effect of Group plotted on standard surfaces. Axial slices provided to show thalamus and ventral striatum regions. **B)** Region-by-region matrix indicating strength of effect of autism versus control. A main effect of Group (Autism vs. Control) was tested on region-by-region correlation levels. Overall, autism participants showed increased correlation between regions during the social interaction tasks, relative to controls (lower triangle unthresholded, upper triangle False-Discovery-Rate corrected to *q*<.05). Increased functional connectivity was correlated with more extensive social impairment measured by the Social Responsiveness Scale (yellow-outlined squares).

We also identified region pairs whose correlation was predicted by the extent of social impairment (indexed by the Social Responsiveness Scale). The most strongly implicated areas were in lingual gyrus bilaterally, the middle cingulate, precuneus, and right inferior frontal gyrus (see yellow squares in Fig. 2B). All results except one were positive correlations, with one region pair showing a negative correlation (higher score −> lower functional connectivity levels between the left cerebellum and the left fusiform/middle temporal gyrus). Three of these social impairment severity effects overlapped with the group effects: that is, there were three region pairs whose correlations were both elevated in autism relative to controls, and also greater in autism participants with greater social impairment. These were the right IFG with right parahippocampal gyrus, right parahippocampal gyrus with right precuneus, and right precuneus with bilateral middle cingulate cortex.

### Group by Task interactions

One region – in the right lateral fusiform gyrus -- showed a difference in whole-brain connectedness by Group and Task (*p*<.0001, FWE corrected to *p*<.05). Using this region as a seed revealed 14 additional regions whose correlations with the seed varied by group and task (Fig. 3; Supplementary Table 6). These additional regions were then aggregated with the right fusiform seed into one set of regions (15 total), and their correlations were analyzed with respect to one another (which had not yet been explicitly tested). In addition to the previously determined relationships between the right fusiform seed and the other 14 regions, three more region pairs that did not include the seed also showed correlations that varied by group and task: left posterior STS with left middle temporal temporal cortex, left middle temporal cortex with right posterior STS, and calcarine gyrus with the right precentral gyrus. For illustration of the nature of these interaction effects, mean Fisher-z transformed correlation coefficients for each participant are plotted in Fig. 3C. Overall, autism participants showed a greater between-region increase in correlation during Conversation compared to Repetition than did controls. We also tested for an effect of extent of social impairment involving the Group-by-Task dependent regions. Only one pair survived False Discovery Rate correction: functional connectivity between right extrastriate and left posterior STS showed a positive correlation with extent of social impairment (Supplementary Fig. 4).

**Figure 3:**
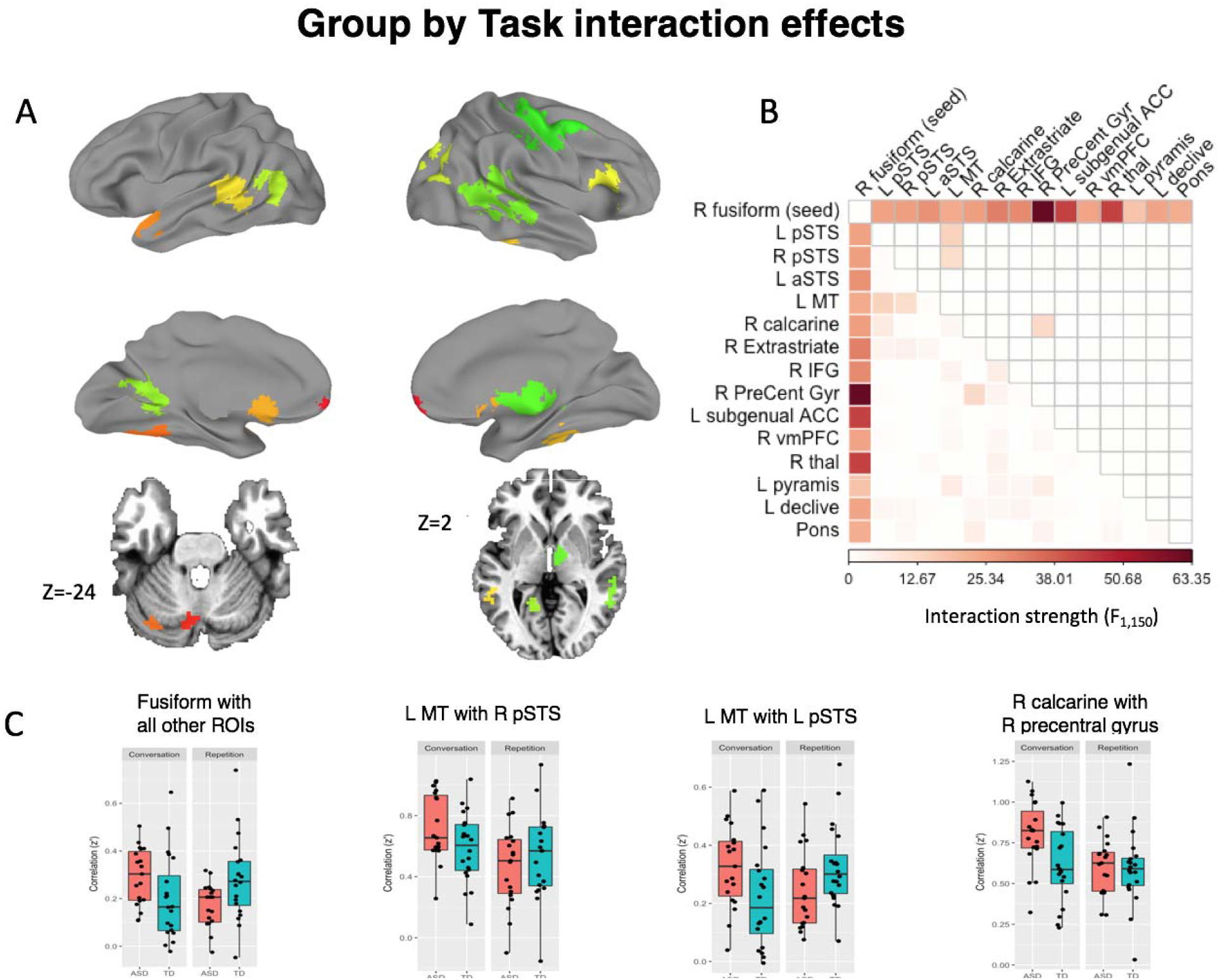
Group by Task interaction effects. **A)** Group-by-task dependent regions of interest. **B)** Region-by-region matrix. Colors indicate larger F-statistics for the interaction of Group*Task. Lower triangle = unthresholded values. Upper triangle indicates correction by False Discovery Rate to *q*<.05. **C)** For significant results, the raw functional connectivity values (Fisher-z transformed correlation coefficients) were extracted and plotted by Group and Task for visualization. Autistic participants showed increased correlation levels during Conversation compared to Repetition. During Conversation, autistic participants showed greater correlation levels between regions than controls.

### Comparison of results during Task and Rest

Both the main effect of Group and the interaction of Group and Task condition (Conversation vs. Repetition) indicated that task-based functional connectivity in autism was greater than in controls (i.e. hyper-correlation). This pattern departs qualitatively from the results reported by some previous data-driven studies of whole-brain functional connectivity in male adolescents and adults, including those by our own lab (e.g. Gotts et al., 2012; Ramot et al., 2017; see also Anderson et al., 2011; Cerliani et al., 2015; Cheng et al., 2015; Di Martino et al., 2014). To examine this potential discrepancy, we therefore compared the patterns of task-based and resting-state functional connectivity using the regions showing a main effect of Group in the current study. The resting-state data have been used in a variety of studies published previously by our lab (Gotts *et al.*, 2012; 2017; Gotts, Jo, *et al.*, 2013; Gotts, Saad, *et al.*, 2013; Plitt *et al.*, 2015; Power *et al.*, 2017; Ramot *et al.*, 2017), with groups matched on age, IQ, head motion, and measures of temporal signal to noise ratio. Region-by-region results are shown in Fig. 4, separately for Task (pooling Conversation and Repetition) and Rest. As already shown above, increases in task-based functional connectivity are observed in autism for these regions (Fig. 4A). Interestingly, increases are also observed during rest, particularly involving the left thalamus (with virtually all of the other regions) and right ventral striatum (with right parahippocampal gyrus, right superior frontal gyrus, and bilateral medial somatosensory cortex, consistent with previous reports by Di Martino *et al*. (2011), Cerliani et al., (2015), and Cheng et al. (2015). These common effects are highlighted as state-invariant, or “core”, region pairs using green squares in Fig. 4C. A number of region pairs also differ significantly by group and state (i.e. Group X State interaction; black squares in Fig. 4C), with many pairs showing increases during Task and lacking significant differences in Rest, and one region pair (left somatomotor with right anterior STS) showing a significant decrease in Rest and lacking differences in Task.

**Figure 4:**
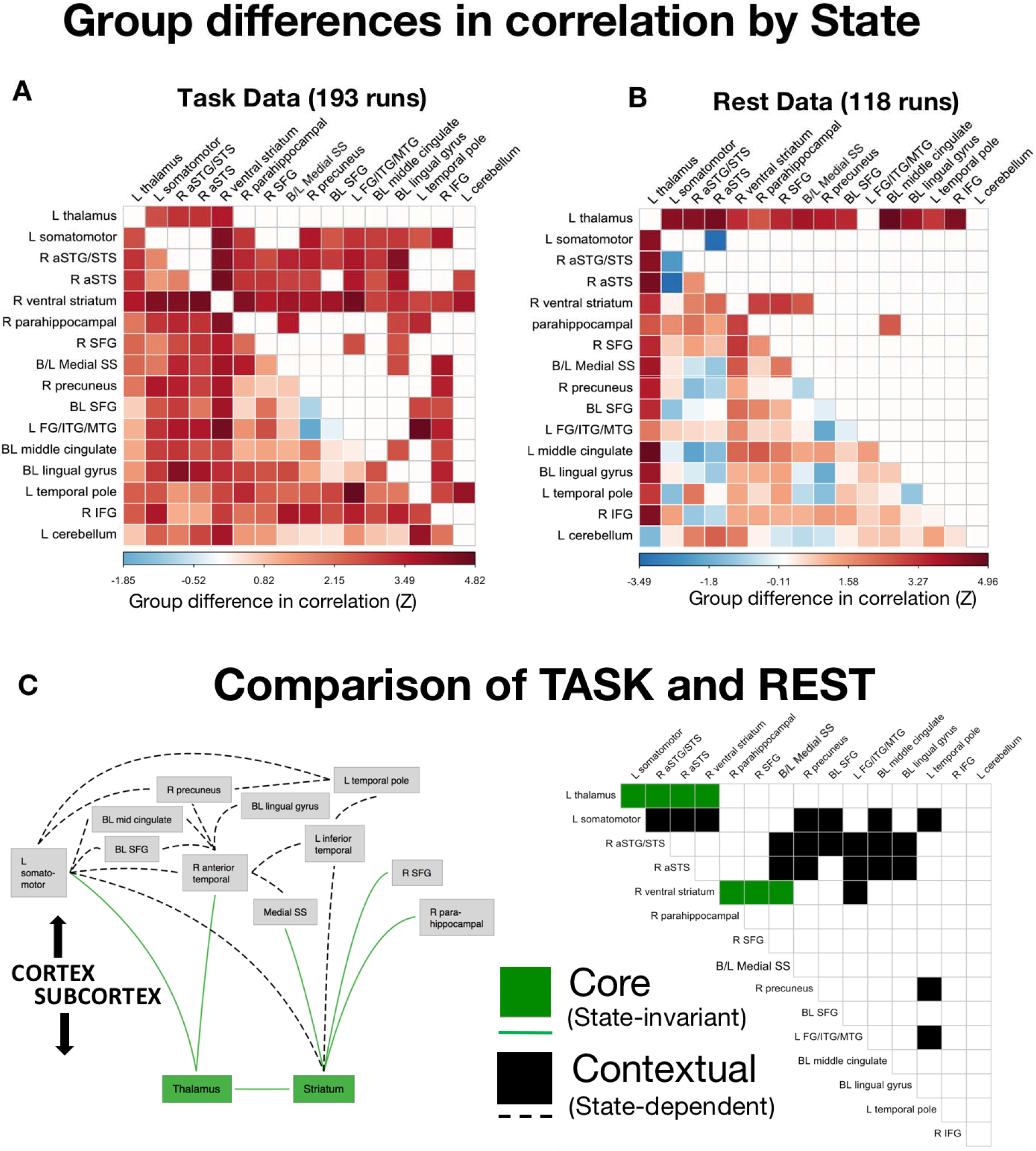
Comparisons of Task and Rest. **A.** Group differences in region-by-region functional connectivity during task in task-defined regions. Higher values indicate larger Autism>Control differences. **B.** Group differences in region-by-region functional connectivity in resting-state data, measured in the same task-defined regions. **C.** Green squares and lines indicate pairs of regions with Autism>Control differences that replicate across Task and Rest. These “core” effects involved the left thalamus and the right ventral striatum (corrected by False Discovery Rate to *q*<.05 in both Task and Rest). Black squares and lines indicate “contextual” effects occurring between cortical areas (significant Group by State interaction, corrected by False Discovery Rate to *q*<.05).

Given the relative lack of differences seen in the Rest data when using the regions detected during Task, we also compared whole-brain connectedness between autism and control at Rest. The results during Rest are shown in Fig. 5 in blue, with the regions showing a main effect of Group during Task shown in red for comparison (voxelwise threshold of *p*<.001 for both Task and Rest, corrected to *p*<.05 by cluster size). Decreases in resting-state functional connectivity in autism are indeed quite prominent, but they overlap very little with the regions showing increases during Task, explaining the lack of decreased functional connectivity seen in Fig. 4B. Despite these state-related differences, the prominent state-invariant results involving the thalamus and ventral striatum (shown in green in Figs. 4C and 5) highlight notable mechanistic commonalities between Task and Rest.

**Figure 5:**
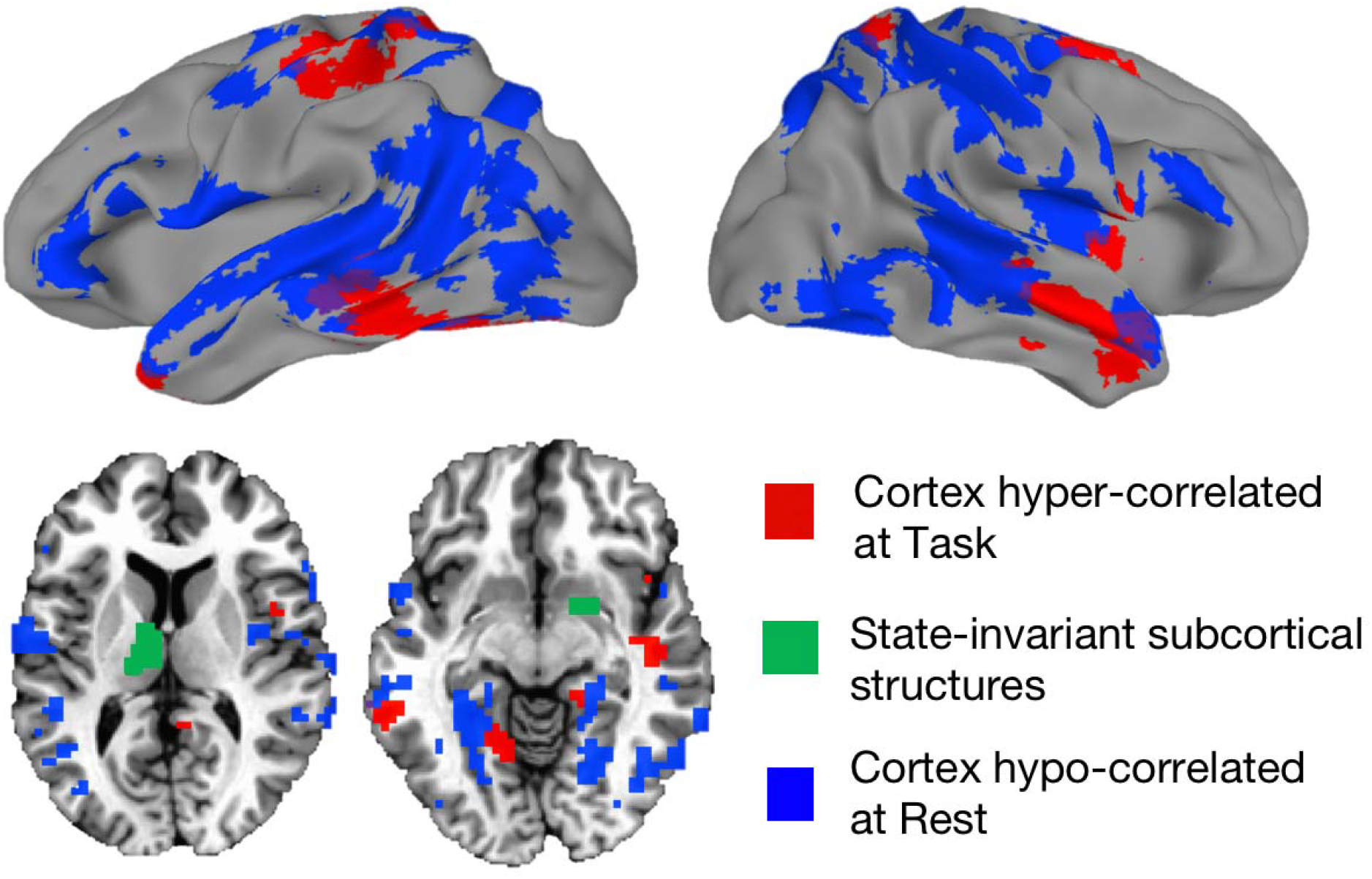
Regions showing the greatest hyper-correlation at Task, and the greatest hypo-correlation at Rest. Autism>Control cortical Task-defined regions (red) shown along with the strongest Control>Autism effects in resting state data (blue). The left thalamus and right ventral striatum regions from the present study are colored green. These subcortical structures showed the greatest state-invariance – with the same differences in correlation with cortex during both task and rest.

## Discussion

The purpose of this study was to characterize neural activity during spontaneous conversation in autism. Participants engaged in face-to-face spoken interactions while being scanned with fMRI. Our design used two groups (people with autism vs. controls) who performed two tasks (Conversation vs. Repetition). Comparing neural activity during the tasks (across groups), greater whole-brain correlation was observed during Conversation than Repetition in regions involved with communication and social processing. Comparing the groups (across tasks), greater task-driven functional connectivity was observed for the autism than the control group, mainly between regions involved with communication and sensorimotor processing. Furthermore, functional connectivity among many region pairs, especially those involving right parahippocampal gyrus, right precuneus, right IFG and bilateral middle cingulate, was strongest in subjects with greater parent-reported social impairments observed outside of the laboratory. Next we examined neural activity differences between groups as a function of task. We found that, particularly among visual and social areas and under high social demand, functional connectivity in the autistic (but not control) participants increased, and increases were related to social impairment. Finally, we examined the relationship between group differences in task-driven functional connectivity and group differences in previously published resting-state connectivity of autism and control participants that instead exhibited a pronounced decrease in cortico-cortical correlations in autism. We found that some region pairs that were hyper-correlated for autism in Task were also hyper-correlated at Rest, and that these pairs involved the thalamus and striatum. By contrast, many more widespread regions showed strong context-specific effects (i.e. Group by State interactions), with regions hyper-correlated during Task failing to show differences in Rest and regions hypo-correlated during Rest failing to show differences in Task (Fig. 4 and 5).

### Increased functional connectivity in autism

We observed widespread increased functional connectivity in autism at Task. These increases were not easily explainable by head motion or other residual global artifacts because measures of these artifacts were covaried in all analyses. This is consistent with at least one other study of functional connectivity during language processing that showed increased occipital recruitment in autism during category judgments of visually-presented nouns (Feelings vs Tools and Colors; (Shen *et al.*, 2012). It also accords with activation studies showing autistic children recruit more areas than control children when judging emotion and irony in prosody (Colich *et al.*, 2012; Eigsti *et al.*, 2012; Wang *et al.*, 2006), or incorporating real-world knowledge in sentence processing (Groen *et al.*, 2010; Tesink *et al.*, 2009). However, other studies of functional connectivity during language processing that required making judgments about imagery (Kana, 2006) or answering questions about agent-patient relationships in visually-presented sentences (Just *et al.*, 2004) have only shown decreased functional connectivity. It may be that tasks with greater sensory demands (e.g. those involving speech vs. mere reading), tasks with social components (e.g. about emotions, sarcastic/ironic prosody, and pragmatics) or tasks requiring sustained attention are more likely to elicit hyper-correlation among cortical areas in people with autism. Moreover, not all of these studies have compared groups with matched behavioral performance (as in the present study).

### Increased correlation during task versus decreases during rest

For adolescent participants with autism, several previous studies of resting-state fMRI functional connectivity have observed decreased correlations among regions engaged in aspects of social processing, such as the STS, medial prefrontal cortex, left IFG, and somatosensory cortex (e.g. Anderson et al., 2011; Cerliani et al., 2015; Cheng et al., 2015; Di Martino et al., 2014; Gotts et al., 2012; von dem Hagen et al., 2013). Some of these decreases are furthermore associated with impairments of social functioning in autistic participants as measured by the Social Responsiveness Scale, which establishes that the functional connectivity decreases are relevant to behavior (Di Martino, Shehzad, *et al.*, 2009; Gotts *et al.*, 2012; Ramot *et al.*, 2017). However, in our current study, correlated inter-regional activity was increased in autism, and these increases were also associated with impairments of social functioning using the same behavioral measure. When we analyzed our previously published resting-state data using the same regions of interest, we found that rather than highlighting a discrepancy between the two datasets in terms of increased versus decreased correlations, a common set of inter-regional relationships held across both sets. These common relationships involved the thalamus and striatum, which were hyper-correlated in autism with each other and with cortical areas. The left thalamus and caudate nucleus exhibited greater correlation with the left somatomotor cortex, the right anterior STS, and the right ventral striatum in autistic participants. The right ventral striatum further exhibited greater correlation in autism with the bilateral medial somatosensory cortex, the right parahippocampal gyrus, and the right superior frontal gyrus. Taken together, the current findings highlight two classes of relationships among regions — “core” state-invariant relations and “contextual” state-dependent relations.

### Context-specificity of cortico-cortical correlations

As discussed, many results were state-dependent – regions that were hypercorrelated during task appeared to function at typical levels or were even hypo-correlated during rest. Futhermore, in our task data, participants with more extensive social impairment showed the greatest inter-regional functional connectivity. Interestingly, an inverse relationship between functional connectivity and social impairment (using the same measure) was found in resting state data, as reported in Gotts et al. (2012) and Ramot et al., (2017). There, participants with greater parent-reported social impairment showed more strongly *decreased* inter-regional correlation. How can this apparent discrepancy be explained? One possibility is that the increased functional connectivity at task is a compensatory neural strategy (c.f. Shen *et al.*, 2012). On this account, more autistic participants with greater social impairment would require greater inter-regional correlation to perform social tasks similarly to controls. This compensatory neural strategy could be deployed in social situations outside the laboratory, too. Indeed, our autistic participants had normal or high IQs and language ability, and their lives seemed in many ways similar to those of controls. Like the control participants, the participants with autism talked (during the experiment) about attending schools, holding jobs, and interacting with peers. In such situations, many simple conversations are likely to occur, most of which are likely to be “successful” in the sense that conversational goals – requesting or providing information, assistance, praise, guidance and so on – are accomplished. Why, then, did our autistic participants receive diagnoses if they are able to compensate and function at nearly normal levels? Even a highly successful neural strategy could result in slight but detectable behavior differences – i.e., the compensation is not perfect, but it is “good enough” to allow some people with milder autism to function (and to make autism diagnosis a non-trivial procedure requiring hours of observation by trained professionals (Lord *et al.*, 2000)). Whether compensatory hyper-correlation observed in this study would generalize to participants with more severe forms of autism is a question that future studies should address.

Notably, the areas that were most severely hypo*-*correlated at Rest and are negatively correlated with degree of social impairment (Fig. 5, blue areas) do not overlap well with the areas that show the highest degree of hyper-correlation during Task (Fig. 5, red areas). This, too, may be consistent with a compensation account. If connectivity among some social brain areas is intrinsically disordered in autism (i.e. Fig. 5 blue areas), perhaps the compensation for these disordered connections takes place via other, spatially non-identical cortical areas (red areas), bringing regions that are hypo-correlated at Rest up to control levels. Novel, compensatory routing among cortical areas may take place via the thalamus, discussed below. Critically, these effects did not appear to manifest in the local activity levels as indexed by a general linear model analysis (Supplementary Materials, Supplementary Fig. 3), helping to establish that these effects are more selective to inter-regional interactions rather than simple activity levels in those regions.

### Core features: abnormal thalamic and striatal interactions – a gating issue?

The thalamus and basal ganglia (within which the striatum resides) are strongly anatomically interconnected with each other and with the entire cerebral cortex. Through these connections, both structures gate transmission between distal cortical regions (McNab and Klingberg, 2008; Sherman, 2007). Could abnormal gating explain the abnormal cortical interactions? In our study the gates were ‘wide open’, especially the intrinsic connections measured during resting state: of the 14 task-defined cortical regions, 12 were intrinsically hyperconnected with the thalamus during rest, even after statistical correction (Fig. 4B). The fact that these regions were defined independently of the rest data is striking and suggests that intrinsic thalamocortical connectivity could partially determine which areas of cortex are likely to exhibit hyper-correlations during sustained or social tasks.

In an early study, Mizuno et al. (2006) observed increased thalamocortical correlation during a non-social visuomotor task (N=8) and suggested that it may be ‘hyperfunctional’, helping to compensate for decreased cortico-cortical connectivity (Mizuno *et al.*, 2006). This is unlikely (and distinct from our current proposal), given that increased thalamocortical correlation is also observed at rest (Cerliani *et al.*, 2015; Cheng et al., 2015; Di Martino *et al.*, 2011; 2014; Nair *et al.*, 2014). Instead, we propose that the context-dependent increases may play this compensatory role. One previous study has examined correlation of brain activity in both task and rest states in autism, but exclusively focused on cortical results. You et al. (2013) found that overall distal connectivity was heightened in autism during a non-social sustained attention task, but decreased at rest (You *et al.*, 2013). Furthermore, they reported that inattention problems in everyday life were associated with a greater increase in functional connectivity from rest to task in autism, from which they conclude that the increased task-driven functional connectivity is maladaptive. However, it is difficult to distinguish maladaptive from compensatory hyper-correlation in the You et al. study as the autism and control groups were not matched for behavioral performance on the in-scanner task.

### Limitations and future directions

Future work should seek to characterize the differences in functional connectivity not just between Task and Rest states, but between different types of tasks. It is almost certain that functional connectivity patterns will vary based on task constraints, and these differences serve as the basis of results in studies that employ techniques such as Psychophysiological Interaction (PPI: Friston et al., 1997) and Dynamic Causal Modeling (DCM: Friston *et al.*, 2003). In the present study, we observed increased functional connectivity in the ASD subjects that strongly implicated right hemisphere regions known to be involved with communication and vocal processing, such as right STS and right IFG. Other tasks that require different sorts of behavior, or with different attentional and social demands, may elicit different patterns. We hope that the comparison between a highly demanding social task and resting state, which we show in this paper, can serve as a starting point for the characterization a broader set of cognitive states.

One caveat concerning the results is that, even when wearing noise-cancelling headphones, the sensory stimulation within an MRI scanner can be intense. A growing literature suggests that autistic individuals may have abnormal sensory and motor processing (Cerliani, *et al.*, 2015; Nebel, *et al.* 2016) and also increased anxiety (see South & Rodgers, 2017 for review). This could potentially give rise to a situation where abnormal connectivity between sensorimotor processing and emotion-related regions is observed in people with autism because of the MRI scanner environment itself. In this regard, it will be important for future studies to re-examine altered dynamics in ASD using quieter neuroimaging methods such as MEG. Along these lines, we have previously noted an encouraging correspondence between our published resting-state fMRI results and MEG phase locking measures (e.g. Ghuman et al., 2017; see Picci et al., for review of recent MEG studies), thus suggesting that at least some of the altered dynamics we report here were not due to intense sensory stimulation.

A limiting factor in the interpretation of our results is that all participants were male and high functioning. The reason for this was to promote consistency with the bulk of prior studies and to limit the mixture of potentially distinct patterns that may occur in females (e.g. Alaerts *et al*., 2017), who are rarer in frequency and more difficult to recruit in sufficient numbers to match those of males. The sample sizes were also more typical of standard fMRI task-based studies (~ 20 participants per group), which is substantially smaller than many of the more recent resting-state studies. We acknowledge that these issues limit the generalizability of the current study, and we hope this may be addressed with future research with larger samples of both males and females, as well as with the inclusion of lower-functioning participants. Another issue concerns the fact that while Repetition and Conversation differ in social demand, they differ in other aspects as well (e.g. mnemonic or linguistic demand). Future work on the neurobiology of social aspects of conversation should seek to refine baseline and control conditions for spontaneous conversation.

To our knowledge this study is the first to characterize brain activity in autism during the context most similar to the gold standard of autism diagnosis (Lord *et al.*, 2000)—face-to-face spoken interactions. It also emphasizes the importance of characterizing autistic brain organization in diverse task contexts and cognitive states. Further research should seek a more mechanistic understanding of whether subcortico-cortical connections may play a causal role in abnormal state-dependent cortico-cortical connections, and how disordered neural connections may be changed via interventions (Ramot *et al.*, 2017).

## Acknowledgments

We thank Bako Orionzi, Ian Eisenberg and Cynthia Peng for their assistance and our participants and their families for their participation.

## Abbreviations

BOLD: Blood Oxygen Level Dependent
IFG: inferior frontal gyrus
IQ: Intelligence Quotient
PCA: principal component analysis
ROI: region of interest
SD: standard deviation
STS: superior temporal sulcus
TR: Repetition Time

